# Thermodynamic and Kinetic Characterization of Protein Conformational Dynamics within a Riemannian Framework

**DOI:** 10.1101/2021.01.19.427358

**Authors:** Curtis Goolsby, Ashkan Fakharzadeh, Mahmoud Moradi

## Abstract

We have formulated a Riemannian framework for describing the geometry of collective variable spaces of biomolecules within the context of molecular dynamics (MD) simulations. The formalism provides a theoretical framework to develop enhanced sampling techniques, path-finding algorithms, and transition rate estimators consistent with a Riemannian treatment of the collective variable space, where the quantities of interest such as the potential of mean force (PMF) and minimum free energy path (MFEP) remain invariant under coordinate transformation. Specific algorithms within this framework are discussed such as the Riemannian umbrella sampling, the Riemannian string method, and a Riemannian-Bayesian estimator of free energy and diffusion constant, which can be used to estimate the transition rate along an MFEP.

## I. INTRODUCTION

Biomolecular simulations have made enormous progress in recent years. The advent of MD in order to study biomolecular phenomena [1–3] has given researchers new insights into previously unviewable phenomena. MD has overcome [4, 5] the common limitation of experimental techniques which forces the researchers to choose between high-resolution static (such as X-ray crystallography) or low-resolution dynamic (such as single-molecule fluorescence resonance energy transfer spectroscopy) pictures of biomolecular systems. MD does however have a few drawbacks. A key issue is the “timescale gap” in that MD simulations typically have a shorter timescale than many relevant biological phenomena. A related issue is the metastability; the system gets trapped in local free energy minima, preventing the system from evenly visiting the entire free energy landscape. Various enhanced sampling techniques and path-finding algorithms have been developed over the last few decades in order to overcome these limitations [6–22]. These techniques are often successful for simple toy models (e.g., dialanine peptide [13, 23–27]); however, practical applications remain quite challenging.

Perhaps the most obvious line of attack for solving the sampling problems is simply to have stronger computing power and more efficient or specialized computer hardware. In the past 40 years, extraordinary advances have been made in this regard. Current state of the art architectures such as very large [28–30] or MD specialized [31, 32] supercomputers and GPU-enabled computing [33–36] have capabilities that allow for simulations with large number of particles to run over long periods of time. The size of supercomputers have also given rise to algorithms developed for speeding up the calculations within brute force MD such as particle mesh Ewald [37] and dynamic load balancing [38]. The focus of this article, however, is the enhanced sampling techniques that are specifically based on enhancing the sampling within a statistical mechanical framework, in particular those methods that, in some form or another, use “collective variables” within their formalism.

In order to obtain relevant thermodynamic and kinetic [39] properties of a statistical mechanical system, one must integrate over high dimensional spaces, which requires large sets of independent and identically distributed samples. In order to manage the dimensionality of biomolecular systems, the assumption is often made that the vast majority of the high-dimensional space is practically “empty” (or occupied by microstates of negligible probability) and the occupied space can be approximated by a lower-dimensional manifold which contains all relevant conformations from stable states to important transition states. Ideally, any elementary reaction, can be thought of as a transition between two stable states, characterized by a transition pathway [40, 41] (or committor function [42]).

The dimensionality reduction may be employed explicitly or implicitly in a sampling scheme. For instance, a path-optimization technique [14, 26, 43–46] can be thought of as a dimensionality reduction technique, if the optimized transition path can be approximated as a thin transition tube, where the areas outside the tube are associated with low probabilities. In this case, one may parametrize this path and potentially deviation from this path [44, 45, 47] and use them as collective variables. Other methods attempt to identify the intrinsic manifold by using statistical learning methods such as principal component analysis [48], isomap [40], and diffusion map [41, 49]. Often these techniques are used to analyze MD trajectories [40, 41, 48, 50] or are combined with enhanced sampling as in metadynamics [51, 52] or adaptive biasing force [53].

Whether a set of collective variables is defined in a systematic manner as described above or it is defined intuitively, it is a convenient way of reducing the dimensionality of the configuration space in both free energy calculation methods and path-finding algorithms. Various algorithms have been developed to estimate free energies [6, 8, 9, 13, 27, 54–59] or find transition pathways [14, 26, 43–46, 60] in collective variable spaces. What is mostly missing is a robust theoretical framework that allows for a rigorous treatment of the issues one needs to deal with when working with collective variables. For instance, the collective variables used in collective-variable based enhanced sampling methods are often nonlinear transformations of atomic coordinates. This complicates their application as noted previously by Johnson and Hummer [61], who show the conventional minimum free energy path (MFEP) obtained from various path-finding algorithms are collective-variable specific and are not invariant under nonlinear coordinate transformations. Such difficulties have often been ignored in the past in the majority of the applications of the collective-variable based simulations; however, there has been attempts in addressing them as in the aforementioned work [61] or within the framework of Transition Path Theory [43, 62]. We recently introduced a Riemannian framework for the rigorous treatment of the collective-variable based spaces within the context of enhanced sampling and path-finding algorithms [63], where the distance, free energy, and MFEP are redefined in a way that are all invariant quantities and do not change under coordinate transformations. Our previous work focused on the thermodynamic characterization of protein dynamics within a Riemannian diffusion model [63]. Here we extend the formalism in two different directions. First, in Section II, we show that the Riemannian formulation is more general than its diffusion model manifestation and applies to any method and formalism that uses collective variables. In Section III, we extend the Riemannian diffusion model by focusing on the kinetics and rate calculation techniques, which we previously did not elaborate on [63].

## II. FROM EUCLIDEAN TO RIEMANNIAN COLLECTIVE VARIABLE SPACE

Euclidean geometry was developed by the Greek Euclid circa 300 BC. Early in the 19th century, mathematicians such as Gauss [64], Schweikart[65], Bolyai [66] and Lobachevsky[67, 68] began to formulate non-Euclidean geometries. With the work of Bernhard Riemann, specifically his lecture “On the Hypotheses which lie at the Bases of Geometry”‘[69], published first in 1873, geometry started exploring new and more diverse applications. Riemann’s work on generalizing the differential surfaces of IR^3^ led to progress in many fields of science. Further progress upon his ideas allowed for the formulation of Einstein’s General Theory of Relativity [70] and progress in group theory [71].

Riemannian geometry provides a robust mathematical framework to develop a formalism for the geometry of collective variable spaces, that are often defined to reduce the dimensionality of the atomic models of macromolecular systems. For instance, consider a transmembrane protein whose transmembrane helices rotate under certain conditions to allow opening or closing of a gate and transporting materials across the membrane. An intuitive collective variable for such a system would be the orientation of the transmembrane helices that can be determined using principal axes of the helices or their orientation quaternions [72, 73]. In order to work in these spaces, we must first have a formalism that allows us to answer common questions which are posed in a typical collective-variable based simulation. Examples of such questions include: what is the distance between two points in the collective variable space? How is the potential of the mean force (PMF) defined at a given point in the collective variable space? How does a biasing potential affect the distribution? How can one find the MFEP? How can we estimate the PMF from biased simulations along the MFEP? How does the system diffuses along a transition pathway? How can we estimate the rate of a transition along a transition pathway? The questions have previously been answered within a Euclidean framework but with our Riemannian treatment of the collective variable space, some of these concepts and quantities need to be revisited. We will begin to answer these questions one at a time.

Imagine a system containing *N* atomic coordinates described by position vector ***x*** under a potential energy surface *V* (***x***). In order to reduce the dimensionality of the system and quantify important functional states, a coarser space is desirable to be defined such that ***xi*** : ℝ^N^ → n, where ***ξ*** is a multi-dimensional collective variable. The PMF is typically defined as:

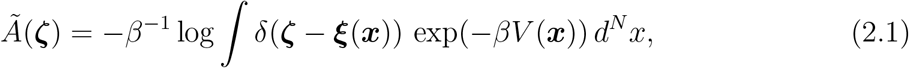

where *Ã*(***ζ***) is the PMF at any given point ***ζ*** in the collective variable space and *δ*(.) is the conventional Dirac delta function.

The PMF as defined above is sometimes considered as the effective potential energy of the reduced system (i.e, a system which is described by the collective variable and not the atomic coordinates). For multi-dimensional collective variable spaces, the MFEP or other related 1D pathways are often used for extensive sampling and characterization in place of the entire multidimensional space. In other words, the multi-dimensional collective variable space can be reduced itself to a one-dimensional curve defined in the collective variable space.

The above approach is quite common and provides a powerful tool for characterizing the energetics of large biomolecular systems. However, we argue here that the above definitions of PMF and MFEP are not well-suited for the purposes that they are defined for. The main problem with these quantities is that they are not invariant under coordinate transformation. Unlike our previous worl [63], the discussions in this Section do not make any assumption regarding the diffusivity of the effective dynamics in the collective variable space and are more general. In the next Section, we will discuss the implications of the diffusivity and the Riemannian diffusion model.

Let us consider a simple toy model to clearly illustrate the non-invariance of conventional PMF. Our toy model is a 1D system in thermal equilibrium with a heat reservoir of temperature *T*, where *β* = (*k*_*B*_*T*)^−1^ = 1. The system is governed by the potential energy *V* (*x*).

The PMF along any collective variable *ξ*(*x*) is defined as:

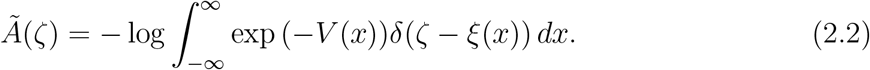

Defining *η*(*ξ*) as the inverse function of *ξ*(*x*) such that *x* = *η*(*ξ*), we can show:

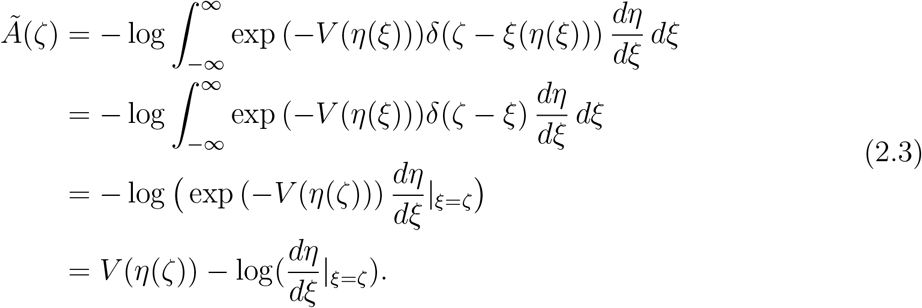

In other words, if 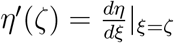, we have:

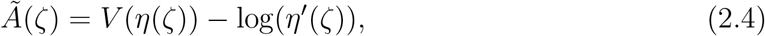

Projecting the collective variable back to the *x* space, we have:

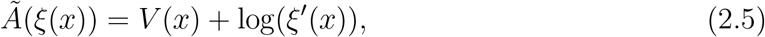

where we have used *ξ*′(*x*) = 1*/η*′(*ζ*) since *η* is the inverse function of *ξ*.

Relations (2.4) and (2.5) clearly show that the shape of PMF is not determined by the potential energy of the system only but it also depends on the derivative of the collective variable. One may use any free energy calculation method (such as umbrella sampling, metadynamics, or ABF) to calculate the PMF; however, without taking into account the second term in Relation (2.4) or (2.5), the PMF’s shape or the shape of its projection onto the *x* space, does not represent the underlying potential energy for such a 1D system.

As an example, let us assume *V* (*x*) = *x*^2^. The shape of PMF may or may not be similar to this potential energy, depending on the definition of collective variable. Figure 1 shows *V* (*x*) along with the PMF as a function of several collective variables (up to an additive constant) that were specifically designed to result in various PMFs. If *ξ*(*x*) = *x*, the PMF would be the same as *V* (*x*). However, if *ξ*(*x*) = *erf* (*x*), the PMF becomes flat since:

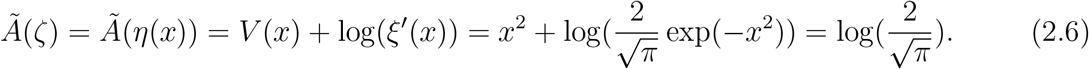

**FIG. 1.**
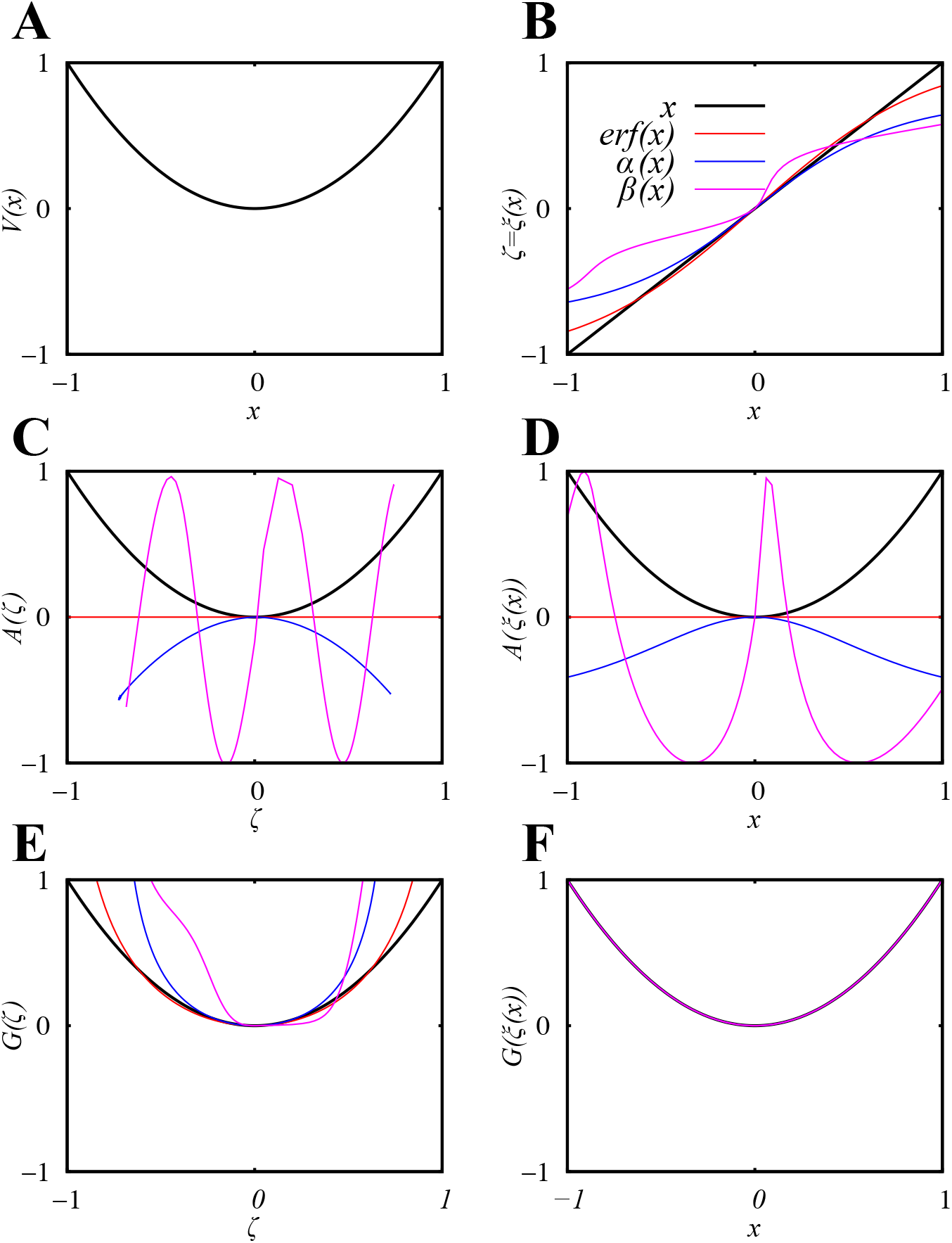
Toy Model: (A) A one-dimensional toy model described by potential energy *V* (*x*) = *x*^2^. (B) Several collective variables (*ξ*(*x*)) are used as examples of smooth coor-dinate transformations 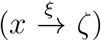. (C) The conventional PMF along the collective variables defined in (B). (D) The projection of conventional PMF (shown in C) onto the *x* space. (E) The Riemannian PMF along the collective variables defined in (B). (F) The projection of Riemannian PMF (shown in E) onto the *x* space (all PMFs are exactly the same when projected onto the *x* space, irrespective of the collective variable used).

Using Relation (2.4), one may solve the following nonlinear differential equation to generate any arbitrary PMF function *Ã*(*ζ*):

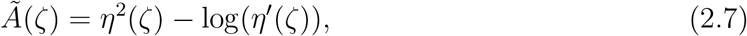

or by some rearrangement:

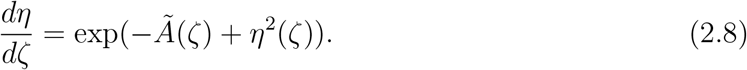

Once the above differential equation is solved for *η*(*ζ*), one can easily find its inverse function *ξ*(*x*). We used this procedure using numerical methods to find functions *α*(*x*) and *β*(*x*) that result in *Ã*(*ζ*) = −*ζ*^2^ and *Ã*(*ζ*) = sin(10*ζ*), respectively. Figure 1 illustrates how the PMF, as defined conventionally based on these collective variables, could qualitatively behave differently from what we expect intuitively from the underlying energetics (here, the potential energy) of this simple system. This is true when PMF is plotted against *ζ* (which is generally expected as *ζ* and *x* are not the same) and more importantly when it is projected back onto the *x* space.

The above example clearly shows that the PMF could be quite misleading if interpreted as a typical potential energy surface, where the minima are considered as locally stable states and the maxima are interpreted as transition states. However, since the collective variable function *ξ*(*x*) is known, log(*ξ*′(*x*)) can also be calculated and subtracted from the PMF to result in *V* (*x*). To do this, first one needs to project *Ã*(*ζ*) onto the *x* space by using *ζ* = *ξ*(*x*) before subtracting log(*ξ*′(*x*)) (Relation (2.5)). Alternatively, one may add log(*η*′(*ζ*)) to *Ã*(*ζ*) to get *V* (*η*(*ζ*)) (Relation (2.4)) and then project that onto *x* to get *V* (*x*). We define an invariant PMF as:

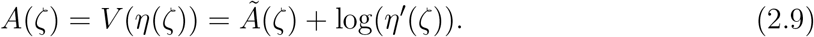

Note that *A*(*ζ*) is a function of *ζ* similar to conventional PMF but *A*(*ξ*(*x*)) = *V* (*x*) for any *ξ*(*x*). We argue *A*(*ζ*) is conceptually more useful than *Ã*(*ζ*) since its connection to *V* (*x*) is more straightforward and its minima and maxima correctly represent the minima and maxima of *V* (*x*). Figure 1 illustrates how the same collective variables used for conventional PMF calculations can also be used to calculate the invariant PMF *A*(*ζ*). The resulting PMFs all qualitatively look similar but once projected back onto the *x* space, they all result in an identical function, i.e., *V* (*x*).

The above definition of invariant PMF can be reformulated as below:

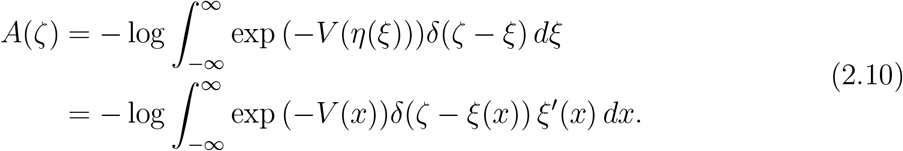

Assuming *x* = *η*(*ζ*) is the only solution to *ζ* = *ξ*(*x*) (i.e., assuming *ξ* is an invertible function of *x*), we can show *δ*(*ζ* − *ξ*(*x*)) = *δ*(*x* − *η*(*ζ*))*/ξ* ′ *x*) (which is the result of identity *δ*(*y*) = *i δ*(*x* − *x*_*i*_)*/y* ′ (*x*_*i*_), where *x*_*i*_’s are the roots of *y*(*x*)). Therefore, using Relation (2.10), we have:

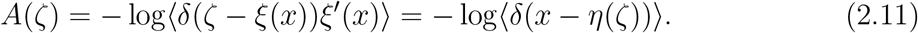

After illustrating the noninvariance nature of conventional PMF and showing that the problem can be resolved by modifying the definition of PMF, we can now generalize the definition of PMF to multidimensional spaces. Relation (2.11) is very easily generalizable for a full transformation from N-dimensional ***x*** to n-dimensional **Ξ**(***x***), where we also no longer assume *β* = 1:

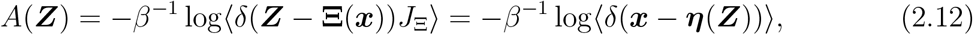

in which *J*_Ξ_ is the determinant of Jacobian ***J***_Ξ_ of transformation **Ξ**. Unfortunately, the full transformation is generally not desirable and ***η***(***Z***) is not available.

If a collective variable ***ξ***(***x***) is lower-dimensional than ***x***, one can assume ***ξ***(***x***) is part of a full transformation. First, we assume *ξ*(***x***) is one-dimensional to simplify the discussion. We can keep the definition of *ξ*(***x***) general, yet choose **Ξ**(***x***) to be the full transformation, such that it contains *ξ*(***x***) and a *N* − 1-dimensional vector orthogonal to *ξ*(***x***), denoted by ***ϕ***(***x***), where orthogonality implies **∇***ξ* · **∇***ϕ*_*i*_ = 0 for all *i*. Here **∇** is the gradient in *x*. Now we can write:

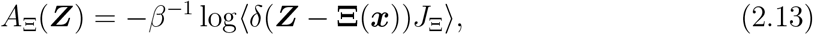

Since the determinant of the product of matrices is equal to the product of their determinants, we can also write:

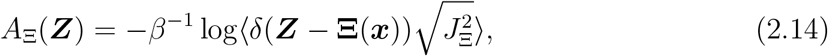

where 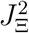 is the determinant of matrix 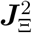. Subsequently, one can write:

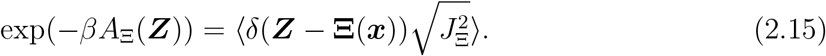

Since *ξ*(***x***) is orthogonal to all *ϕ*_*i*_(***x***), one can easily show that 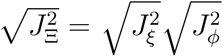, where 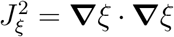 and 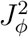 is the determinant of a (*N* − 1) × (*N* − 1) matrix containing elements**∇***ϕ*_*i*_ · **∇***ϕ*_*j*_. We can write:

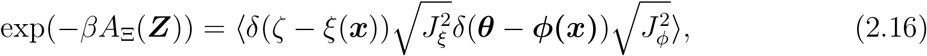

where ***Z*** = (*ζ*, ***θ***). The *N* -dimensional invariant PMF can be used to define the 1D invariant PMF *A*(*ζ*) as:

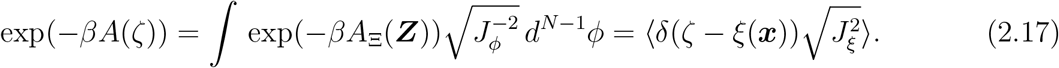

*A*(*ζ*) is invariant since (1) *A*_Ξ_(*ζ*) is invariant and (2) *A*(*ζ*) does not depend on the choice of ***ϕ*** as long as ***ϕ*** is orthogonal to *ξ*. Now we define what we refer to as the metric *g*, using the conditional ensemble average of 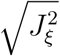:

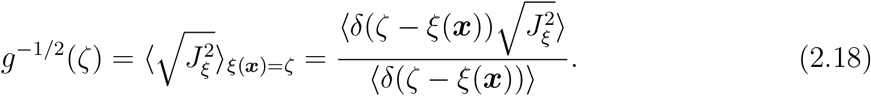

With the above definition of metric, one can now write:

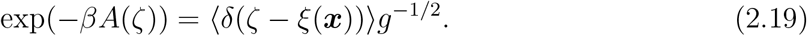

This can be rewritten as:

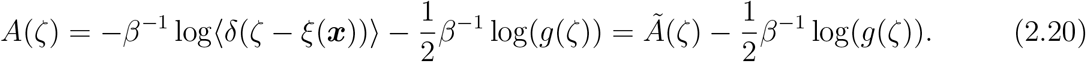

The above definition of 1D invariant PMF *A*(*ζ*) can be generalized to any arbitrary number of dimensions:

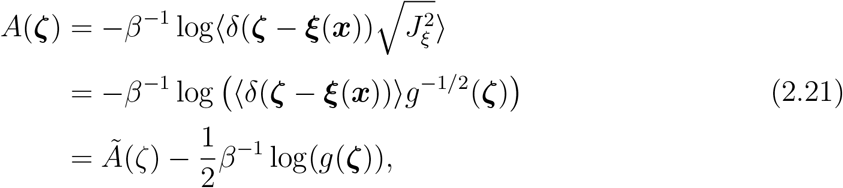

where matrix 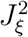 is composed of components **∇***ξ*_*i*_ · **∇***ξ*_*j*_. The metric inverse ***g*** is defined to satisfy the relationship 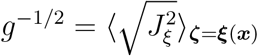. This can be achieved by constructing elements of metric inverse ***g***^-1^ as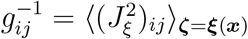.

The above definition of invariant PMF differs from the conventional definition of PMF as it involves the metric tensor ***g***. The metric tensor is a well-known quantity in differential geometry and is needed to do invariant measurements in non-Euclidean spaces. The PMF is only an example of a quantity that can be redefined to be invariant with the help of the metric tensor. The noninvariance of the conventional PMF and any other geometric quantity in the collective variable space is known, although it is often overlooked. The PMF can be made invariant by adding an additional term, which depends on the derivatives of collective variables, as derived above. The difference between the invariant PMF and conventional PMF is related to the Fixman potential as also discussed elsewhere [74, 75]. Note that the Fixman potential was originally developed to relate the PMF associated with a constrained dynamics to that of an unconstrained one [76, 77]. A conceptually more straightforward approach to ensure the invariance of not only the PMF but also any other quantity of interest is to treat the collective variable space as a Riemannian space. In this approach, the metric tensor is used to calculate quantities such as distances, gradients, and integrals in an invariant manner. For instance, for integration, the regular volume element *d*^*n*^*ξ* is replaced by the Riemannian volume element 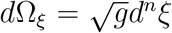, which is invariant under coordinate transformation. Also 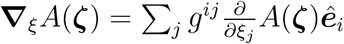 is the invariant gradient, where ***ê***_*i*_’s are unit vectors along *ξ*_*i*_’s.

Riemannian geometry provides the conceptual framework and mathematical tools to treat the nonlinear behavior of curved spaces that are smooth but have potentially different curvatures at different points of space. The rigorous treatment of the collective variable space fits well within the Riemannian geometry framework. In this framework, the Riemannian PMF is defined exactly the same way as the conventional PMF is defined but the Riemannian Dirac delta function replaces the conventional Dirac delta function:

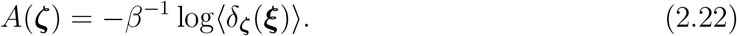

in which *δ*_***ζ***_(***ξ***) is the Riemannian Dirac delta function. Comparing Relation (2.22) to Relation (2.21), one can make a connection between the Riemannian and conventional Dirac delta functions: 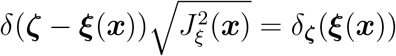.

With the same argument made above, one can show the conventional MFEP is not invariant under coordinate transformation as shown by Johnson and Hummer [61]. Our Riemannian treatment of the collective variable space, however, provides a robust framework for defining the MFEP in an invariant way [63]. The MFEP in a Euclidean space is defined as a path parallel to the gradient of free energy 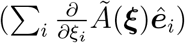. Unfortunately, the conventional MFEP is not invariant under coordinate transformation, which questions its importance as a meaningful quantity. The Riemannian framework allows us to define the MFEP simply based on the Riemannian/invariant gradient of Riemannian/invariant PMF. In other words, not only both PMF and its gradient are well-defined, invariant quantities within the Riemannian framework, the Riemannian MFEP that is simply a path parallel to the gradient of PMF is also well-defined and invariant under coordinate transformation.

## III. RIEMANNIAN DIFFUSION AND TRANSITION RATE ESTIMATION

In the previous Section, we did not make any assumptions regarding the dynamics. However, quantities such as PMF and particularly MFEP are difficult to interpret if the reduced system follows a non-diffusive dynamics in the collective variable space. The intuitive interpretation of minima and saddle points of PMF representing the stable and transition states and the MFEP representing the most probable pathway relies on the diffusive nature of the effective dynamics. Such a condition is only satisfied if at all with specific choices of collective variables. Here, the focus of our discussion is not on how to find such collective variables. However, if we can successfully identify a set of collective variables such that the projected motion of the system on this reduced collective variable space is diffusive, we can write:

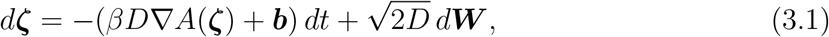

where *D* is the diffusion constant and 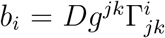, in which 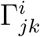’s are Christoffel symbols and Einstein summation convention is used. Note that *D* is not assumed to be position-dependent here. Instead the position dependence is absorbed in metric tensor ***g***. *d****W*** is a Riemannian Wiener process, where 〈*W*^*i*^(*t*)〉 = 0 and 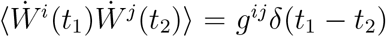, where 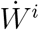 is the time derivative of *W*^*i*^.

Rewriting this equation as Fokker-Planck or Smoluchowski equations [78], we have:

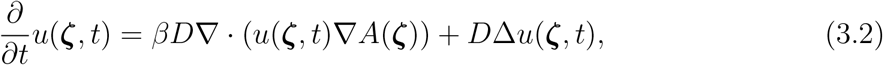

which contains the Laplace-Beltrami operator, 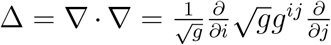, and *u*(***ζ***, *t*) is the probability of finding the system at ***ζ*** at time *t* with a boundary condition of *u*(***ζ***, 0) = *u*_0_(***ζ***).

This summarizes our Riemannian diffusion model, which was previously introduced in Ref. [63]. In the following we derive a new relation that allows the estimation of the PMF, metric, diffusion constant, and transition rate from unbiased simulations performed along an approximate MFEP.

We have previously discussed how one can find a Riemannian MFEP using the Riemannian implementation of string method with swarms of trajectories [63]. Let us assume we have found such a pathway, ***ξ***(*s*), parametrized by its arclength *s*. At any given point along this path, *ê*_*s*_ is the unit vector parallel to ***ξ***(*s*). An *n* − 1 dimensional submanifold (Σ_*s*_) can be defined perpendicular to *ê*_*s*_. Let us assume local coordinates ***ζ*** = (*s*, ***κ***) describes any point on Σ_*s*_. One can write:

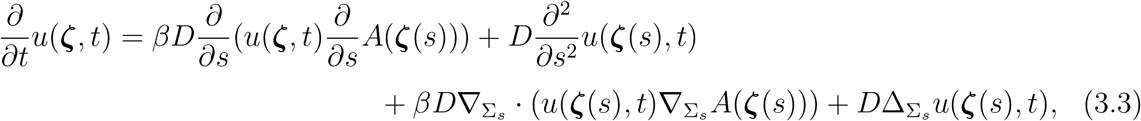

where 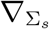 denotes the ∇ operator in the submanifold Σ_*s*_.

Let us now define 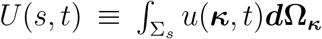, where Σ_*s*_ is an arbotrary portion of the subspace of ***κ*** around its origin. On the MFEP, let us also define a univariate PMF *G*(*s*) = *A*(*ξ*(*s*)).

Continuing, we can integrate over individual terms in Relation (3.3), where the LHS of the equation becomes 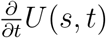 and the RHS terms can be approximated as follows. Assuming *u*(***ξ***, *t*) is much larger within a relatively narrow “tube” around the MFEP, we can integrate over the cross section of this tube and the areas around the tube. In other words, we choose Σ_*s*_ to be a portion of the ***κ*** space that covers the cross section of transition tube that falls within this space as well as some low-probability areas around it. The first term of the RHS of Relation (3.3) will thus be:

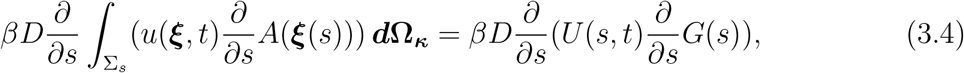

where we define a univariate PMF *G*(*s*) = *A*(*ξ*(*s*)) and assume *A*(*ξ*(*s*)) is more or less constant on any cross section of transition tube perpendicular to the MFEP. The next term becomes 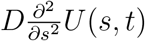 after integration and the third term will vanish assuming 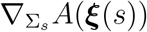 stays close to zero within the tube. To be more precise, we assume the following integral is negligible as 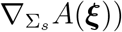 is negligible close the MFEP:

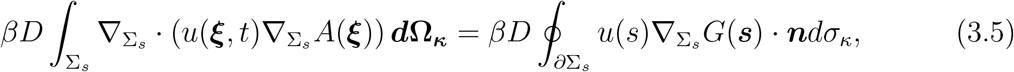

where the divergence theorem is used. Similarly, we can use the divergence theorem to reduce the last term to:

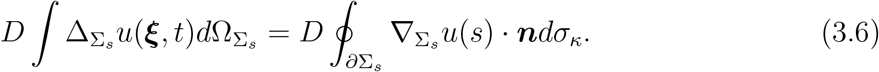

We assume this term is also negligible since the 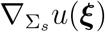 · ***n*** term is only evaluated outside the transition tube and is expected to be close to zero on average. These approximations reduce Relation (3.3) to a one-dimensional diffusion equation in terms of probability density *U* (*s, t*) and the univariate PMF (or potential energy) *G*(*s*):

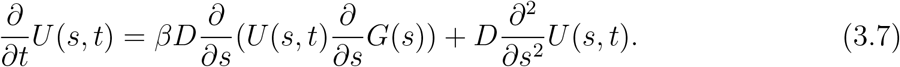

The thin transition tube, used to simplify Relation (3.3) is a common assumption made for path-finding algorithms such as string method [79]. At this point, we have reduced our atomic model, first to a coarse variable space, and then we have focused upon a single dimension, namely the transition pathway of interest.

Relation (3.7) is quite similar to the conventional Smoluchovsky equation in a one-dimensional Euclidean space. However, this is due to the fact that we assumed *s* to represent the geodesic distance along the MFEP path. A slightly more general relation can be derived based on (3.7) for an arbitrary parameter *r*, parametrizing the MFEP path. The *r* space will then be associated with a 1D metric *h*(*r*). We have *ds*^2^ = *h*(*r*)*dr*^2^ or 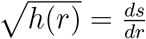. The Riemannian Smoluchowsky equation in then locally written in terms of *r* as:

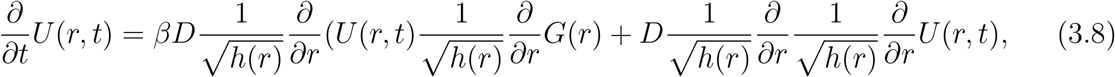

which is slighhtly different from a conventional 1D position-dependent diffusion equation. Here *D/h*(*r*) is equivalent to the conventional 1D position-dependent diffusion constant and *G*(*r*) is the same as the conventional 1D PMF in terms of *r* with an extra term 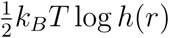, which makes the Riemannian PMF independent of the choice of *r*.

Finally, we can rewrite Relation (3.8) as:

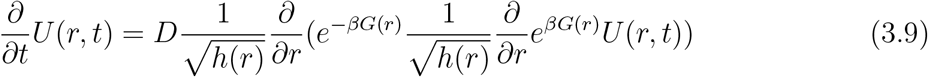

Now suppose that one has been able to identify the MFEP without necessarily quantifying the metric. Relation (3.9) provides a framework to determine the dynamics of the system as long as it stays close to the MFEP transion tube. The PMF and metric along *r* (*G*(*r*) and *h*(*r*)) fully describe the diffusive dynamics and the arbitrary diffusion constant *D* simply determines the unit of *h*(*r*). In Section V, we will discuss in more detail how the estimation of *G*(*r*) and/or *h*(*r*) is possible using unbiased or biased simulations; however, none of these discussions would be relevant without identifying the MFEP.

### Determining the Rate of Transition

Our final formulation mathematically follows closely the work of Hummer [80], except we are working within a Riemannian geometry. We define 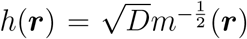 and *H*(***r***, *t*) = *e*^*βG*(***r***)^*U* (***r***, *t*) such that (14) can be rewritten in an easily discretized form:

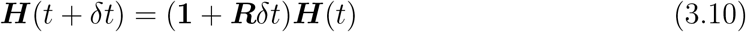

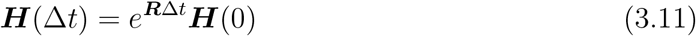

***H*** is here a discretized representation of the transition probabilities which can be determined empirically and R is a tridiagonal matrix for which more can be seen in section 4.

We can self consistently [81] solve by maximizing the likelihood between states at t and t±1 defined by:

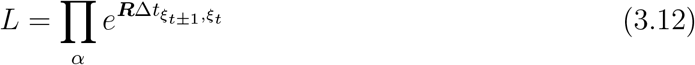

using the tridiagonal ***R*** from (16):

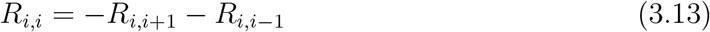

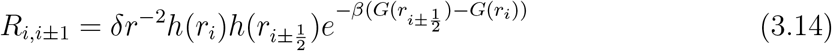

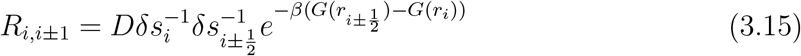

using the relation that 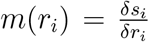 and r values chosen such that *δr*_*i*_ = *δr*_*i*+1_. Now it can be seen that working in a Riemannian collective variable space is valid for the determination of quantities of interest such as rate, free energy calculations, diffusion, and reaction pathways.

## Acknowledgement

This research is supported by the National Science Foundation under grant CHE-1945465.

## References

[1] T. Hansson, C. Oostenbrink, and W. F. van Gunsteren, Current Opinion in Structural Biology 12, 190 (2002).

[2] M. Karplus and J. A. McCammon, Nature Structural Biology 265, 654 (2002).

[3] K. Lindorff-Larsen, S. Piana, R. O. Dror, and D. E. Shaw, Science 334, 517 (2011).

[4] M. Karplus and J. Kuriyan, Proceedings of the National Academy of Sciences, USA 102, 6679 (2005).

[5] J. L. Klepeis, K. Lindorff-Larsen, R. O. Dror, and D. E. Shaw, Current Opinion in Structural Biology 19, 120 (2009).

[6] G. M. Torrie and J. P. Valleau, Journal of Chemical Physics 23, 187 (1977).

[7] S. H. Northrup, M. R. Pear, C. Y. Lee, J. A. McCammon, and M. Karplus, Proceedings of the National Academy of Sciences, USA 79, 4035 (1982).

[8] S. Izrailev, S. Stepaniants, M. Balsera, Y. Oono, and K. Schulten, Biophysical Journal 72, 1568 (1997).

[9] J. Schlitter, M. Engels, P. Krüger, E. Jacoby, and A. Wollmer, Molecular Simulation 10, 291 (1993).

[10] Y. Sugita and Y. Okamoto, Chemical Physics Letters 314, 141 (1999).

[11] Y. Sugita, A. Kitao, and Y. Okamoto, Journal of Chemical Physics 113, 6042 (2000).

[12] D. M. Zuckerman and T. B. Woolf, Physical Review E 63, 016702 (2000).

[13] A. Laio and M. Parrinello, Proceedings of the National Academy of Sciences, USA 99, 12562 (2002).

[14] W. Ren, E. Vanden-Eijnden, P. Maragakis, and W. E, Journal of Chemical Physics 123, 134109 (2005).

[15] E. Darve, D. Rodríguez-Gómez, and A. Pohorille, Journal of Chemical Physics 128, 144120 (2008).

[16] I. Abrahama, S. Jainb, C.-P. Wu, M. A. Khanfar, Y. Kuang, C.-L. Dai, Z. Shi, X. Chen, L. Fu, S. V. Ambudkar, K. E. Sayed, and Z.-S. Chen, Biochemical Pharmacology 80, 1497 (2010).

[17] A. Mitsutake, Y. Mori, and Y. Okamoto, in Biomolecular Simulations, Methods in Molecular Biology, Vol. 924, edited by L. Monticelli and E. Salonen (Humana Press, New York, 2013) pp. 153–195.

[18] M. Moradi, V. Babin, C. Sagui, and C. Roland, in Biomolecular Simulations, Methods in Molecular Biology, Vol. 924, edited by L. Monticelli and E. Salonen (Humana Press, New York, 2013) pp. 313–337.

[19] C. Abrams and G. Bussi, Entropy 16, 163 (2013).

[20] C. Templeton, S.-H. Chen, A. Fathizadeh, and R. Elber, Journal of Chemical Physics 147, 152718 (2017).

[21] L. T. Chong, A. S. Saglam, and D. M. Zuckerman, Current Opinion in Structural Biology 43, 88 (2017).

[22] A. Laio, A. Z. Panagiotopoulos, and D. M. Zuckerman, Journal of Chemical Physics 149, 072001 (2018), https://doi.org/10.1063/1.5049669.

[23] M. Mezei, Molecular Simulation 3, 301 (1989).

[24] R. Crehuet and M. J. Field, Journal of Chemical Physics 118, 9563 (2003).

[25] H. Jang and T. B. Woolf, Journal of Computational Chemistry 27, 1136 (2006).

[26] A. C. Pan, D. Sezer, and B. Roux, Journal of Physical Chemistry B 112, 3432 (2008).

[27] A. L. Ferguson, A. Z. Panagiotopoulos, P. G. Debenedetti, and I. G. Kevrekidis, Journal of Chemical Physics 134, 135103 (2011).

[28] F. Allen, G. Almasi, W. Andreoni, D. Beece, B. J. Berne, A. Bright, J. Brunheroto, C. Cascaval, J. Castanos, P. Coteus, P. Crumley, A. Curioni, M. Denneau, W. Donath, M. Eleftheriou, B. Fitch, B. Fleischer, C. J. Georgiou, R. Germain, M. Giampapa, D. Gresh, M. Gupta, R. Haring, H. Ho, P. Hochschild, S. Hummel, T. Jonas, D. Lieber, G. Martyna, K. Maturu, J. Moreira, D. Newns, M. Newton, R. Philhower, T. Picunko, J. Pitera, M. Pitman, R. Rand, A. Royyuru, V. Salapura, A. Sanomiya, R. Shah, Y. Sham, S. Singh, M. Snir, F. Suits, R. Swetz, W. C. Swope, N. Vishnumurthy, T. J. C. Ward, H. Warren, and R. Zhou, IBM Systems Journal 40, 310 (2001).

[29] T. Diede, C. F. Hagenmaier, G. S. Miranker, J. J. Rubinstein, and W. Worley, Computer 21, 13 (1988).

[30] A. Dubrow, Computing in Science & Engineering 17, 83 (2015).

[31] D. E. Shaw, J. Grossman, J. A. Bank, B. Batson, J. A. Butts, J. C. Chao, M. M. Deneroff, R. O. Dror, A. Even, C. H. Fenton, et al., in Proceedings of the International Conference for High Performance Computing, Networking, Storage and Analysis (IEEE Press, 2014) pp. 41–53.

[32] D. E. Shaw, R. O. Dror, J. K. Salmon, J. P. Grossman, K. M. Mackenzie, J. A. Bank, C. Young, M. M. Deneroff, B. Batson, K. J. Bowers, E. Chow, M. P. Eastwood, D. J. Ierardi, J. L. Klepeis, J. S. Kuskin, R. H. Larson, K. Lindorff-Larsen, P. Maragakis, M. A. Moraes, S. Piana, Y. Shan, and B. Towles, in Proceedings of the Conference on High Performance Computing Networking, Storage and Analysis, SC ‘09 (ACM, New York, NY, USA, 2009) pp. 39:1–39:11.

[33] J. E. Stone, J. C. Phillips, P. L. Freddolino, D. J. Hardy, L. G. Trabuco, and K. Schulten, Journal of Computational Chemistry 28, 2618 (2007).

[34] M. J. Harvey, G. Giupponi, and G. D. Fabritiis, Journal of Chemical Theory and Computation 5, 1632 (2009).

[35] D. E. Tanner, J. C. Phillips, and K. Schulten, Journal of Chemical Theory and Computation 8, 2521 (2012).

[36] A. W. Gotz, M. J. Williamson, D. Xu, D. Poole, S. Le Grand, and R. C. Walker, Journal of chemical theory and computation 8, 1542 (2012).

[37] P. F. Batcho, D. A. Case, and T. Schlick, J. Chem. Phys. 115, 4003 (2001).

[38] J.-L. Fattebert, D. F. Richards, and J. N. Glosli, Computer Physics Communications 183, 2608 (2012).

[39] M. Held and F. Noé, European journal of cell biology 91, 357 (2012).

[40] P. Das, M. Moll, H. Stamati, L. E. Kavraki, and C. Clementi, Proceedings of the National Academy of Sciences, USA 103, 9887 (2006).

[41] A. L. Ferguson, A. Z. Panagiotopoulos, P. G. Debenedetti, and I. G. Kevrekidis, Proceedings of the National Academy of Sciences, USA 107, 13597 (2010).

[42] W. E W. Ren, and E. Vanden-Eijnden, Chemical Physics Letters 413, 242 (2005).

[43] L. Maragliano, A. Fischer, E. Vanden-Eijnden, and G. Ciccotti, Journal of Chemical Physics 125, 024106 (2006).

[44] M. Chen and W. Yang, J. Comput. Chem. 30, 1649 (2009).

[45] G. Díaz Leines and B. Ensing, Physical Review Letters 109, 020601 (2012).

[46] L. Cao, C. Lv, and W. Yang, J. Chem. Theory Comput. 9, 3756 (2013).

[47] D. Branduardi, F. L. Gervasio, and M. Parrinello, Journal of Chemical Physics 126, 054103 (2007).

[48] A. Amadei, A. B. M. Linnsen, and H. J. C. Berendsen, PROTEINS: Structure, Function, and Genetics 17, 412 (1993).

[49] A. L. Ferguson, A. Z. Panagiotopoulos, I. G. Kevrekidis, and P. G. Debenedetti, Chemical Physics Letters 509, 1 (2011).

[50] J. R. Perilla and T. B. Woolf, Journal of Chemical Physics 136, 164101 (2012).

[51] V. Spiwok, P. Lipovová, and B. Králová, Journal of Physical Chemistry B 111, 3073 (2007).

[52] G. A. Tribello, M. Ceriotti, and M. Parrinello, Proceedings of the National Academy of Sciences, USA 109, 5196 (2012).

[53] B. Hashemian, D. Millán, and M. Arroyo, Journal of Chemical Physics 139, 214101 (2013).

[54] T. Huber, A. E. Torda, and W. F. van Gunsteren, J. Comput.-Aided Mol. Des. 8, 695 (1994).

[55] G. Hummer and I. G. Kevrekidis, Journal of Chemical Physics 118, 10762 (2003).

[56] V. Babin, C. Roland, and C. Sagui, Journal of Chemical Physics 128, 134101 (2008).

[57] P. R. L. Markwick, L. C. T. Pierce, D. B. Goodin, and J. A. McCammon, Journal of Physical Chemistry Letters 2, 158 (2011).

[58] T. Hayami, K. Kasahara, H. Nakamura, and J. Higo, Journal of computational chemistry (2018).

[59] F. Paul, Markov state modeling of binding and conformational changes of proteins, Ph.D. thesis, Universität Potsdam Potsdam (2017).

[60] Y. Miao, F. Feixas, C. Eun, and J. A. McCammon, Journal of computational chemistry 36, 1536 (2015).

[61] M. E. Johnson and G. Hummer, Journal of Physical Chemistry B 116, 8573 (2012).

[62] W. E and E. Vanden-Eijnden, Annual Review of Physical Chemistry 61, 391 (2010).

[63] A. Fakharzadeh and M. Moradi, Journal of Physical Chemistry Letters 7, 4980 (2016).

[64] J. Grey (Springer, 2006) Chap. 2.

[65] F. Schweikart and C. Gabler, Die Theorie der Parallellinien, nebst dem Vorschlage ihrer Verbannung aus der Geometrie, Die Theorie der Parallellinien, nebst dem Vorschlage ihrer Verbannung aus der Geometrie (bey Christian Ernst Gabler, 1807).

[66] J. Bolyai, The Science Absolute of Space: Independent of the Truth or Falsity of Euclid’s Axiom XI (which can never be decided a priori) (The Neomon, 1896).

[67] N. I. Lobachevsky and V. Kagan, Geometrical Investigations on the theory of parallel lines; On the foundations of geometry, Vol. 1 (1829–30).

[68] N. I. Lobachevsky and V. Kagan, New Foundations of Geometry with a Complete Theory of Parallels, Vol. 2 (1835-38).

[69] B. Riemann and H. Weyl, in Uber die Hypothesen, welche der Geometrie zu Grunde liegen (Springer Berlin Heidelberg, Berlin, Heidelberg, 1921) pp. 1–47.

[70] A. Einstein, Sitzungsber. Preuss. Akad. Wiss. Berlin (Math. Phys.), 1030 (1914).

[71] J. Milnor, Advances in Mathematics 25 (1977).

[72] G. Fiorin, M. L. Klein, and J. Hénin, Molecular Physics 111, 3345 (2013).

[73] M. Moradi and E. Tajkhorshid, Journal of Physical Chemistry Letters 4, 1882 (2013).

[74] C. Hartmann and C. Schtte, Physica D: Nonlinear Phenomena 228, 59 (2007).

[75] C. Hartmann, J. C. Latorre, and G. Ciccotti, The European Physical Journal Special Topics 200, 73 (2011).

[76] M. Fixman, Proceedings of the National Academy of Sciences 71, 3050 (1974).

[77] E. A. Carter, G. Ciccotti, J. T. Hynes, and R. Kapral, Chemical Physics Letters 156, 472 (1989).

[78] M. v. S. Einstein, A., Untersuchungen uber die Theorie der Brownschen Bewegung, Ostwalds Klassiker der exakten Wissenschaften (Deutsch, 1997).

[79] W. E. and E. Vanden-Eijnden, Journal of Statistical Physics 123, 503 (2006).

[80] G. Hummer, New Journal of Physics 7, 34 (2005).

[81] A. Singharoy, C. Chipot, M. Moradi, and K. Schulten, Journal of the American Chemical Society 139, 293 (2016).

